# Estimation of the transmission dynamics of H5N1 HPAI outbreak in a dairy herd using a modeling approach

**DOI:** 10.64898/2026.09.22.753543

**Authors:** Pranav S. Kulkarni, Sharif S. Aly, Deniece R. Williams, Wagdy R. ElAshmawy, Pranav S. Pandit

## Abstract

The emergence of Highly Pathogenic Avian Influenza (HPAI 2.3.4.4b) in dairy herds in 2024 across 19 states in the United States of America has raised concerns regarding the potential national and global zoonotic impact. All recent modeling efforts implemented homogenous cattle-to-cattle (both intra and inter-herd) transmission models which did not capture the real-world heterogeneity in mixing of animals, individual variations in susceptibility and infectiousness and clinical incidences across pens and lactation groups. The aim of this study was to develop a heterogenous transmission model to estimate the epidemiological parameters for intra-herd HPAI transmission on Californian dairies. We developed a validated stochastic agent-based model to estimate the epidemiological parameters for intra-herd HPAI transmission on California dairies. The hierarchical agent-based model also parameterized stochastic cattle movements within-herd to simulate real dairy management practices. A cow-level SEIR transmission approach was assumed during the outbreak. A novel Bayesian Optimizer with Gaussian Process (BO-GP) was fitted to the agent-based model for validation which converged within 25-40 iterations (out of 100 per farm) with minimal loss over two distinct error metrics, namely, Poisson loss function and temporal distance metric. Our optimized simulations estimated an average R_0_ was around 10.7-10.8 across all farms within the first 15 days of observed outbreak on four dairy farms with a mean effective transmission rate of 4.5% per contact between susceptible and infectious cows within each pen. Our model demonstrated that the movement of cows between pens ensured localized clusters of outbreaks within the sub-herds (pen population) that prolonged the overall outbreak within farms. We estimated the total duration of infection between 14.5 and 28 days, which is higher than the estimates from the homogenous models. With an integrated hierarchical agent-based model combined with Bayesian approximation, we produced actionable insights on the epidemiology of intra-farm spread of HPAI within cow herds, thereby guiding both future model development and applied disease control strategy.

## INTRODUCTION

The emergence of Highly Pathogenic Avian Influenza (HPAI 2.3.4.4b) in dairy herds in 2024 across 19 states in United States of America has raised concerns regarding the potential national and global impact. These concerns were further substantiated by evidence of cattle-to-human transmission during the 2024 outbreak in US dairy herds (Uyeki et al., 2024; Bartlett et al., 2025). Evidence further suggested a single spillover event from wild birds to cattle, resulting in a rapid simultaneous point source epidemic (Nguyen et al., 2025; Rodriguez et al., 2025). The virus disseminated across the US through symptomatic and asymptomatic cattle movement, introducing the virus to dairy populations of other states, and subsequently back into other host species (Nguyen et al., 2025).

Though milking parlors and calving areas were strongly suspected as potential hotspots, it was unclear how they contributed to cattle-to-cattle transmission (Sage et al., 2024; Stenkamp-Strahm et al., 2025). Insights regarding intra-farm dynamics and spread of infection would pave the path for implementing biosecurity measures and mitigation strategies that would ultimately reduce the potential impact of subsequent outbreaks (Rodriguez et al., 2024; Kamel et al., 2025). Quantifying intra-farm cattle-to-cattle transmission was crucial to reveal the early dynamics of disease spread, to inform on-farm management, assessment of further HPAI risk for dairy herds and estimating epidemiological parameters such as transmission rate and basic reproduction number (Malladi et al., 2026).

However, the current modeling efforts implemented homogenous cattle-to-cattle (both intra and inter-herd) transmission models which did not capture the real-world heterogeneity in mixing of animals, individual variations in susceptibility and infectiousness and clinical incidences across pens and lactation groups (Bellotti et al., 2024; Rawson et al., 2025; Malladi et al., 2026). Intra-herd transmission is also expected to be affected by direct interactions through shared milking machines, cross-parenting, influx of pregnant springers in open herds, as well as indirect interactions through occupying the same pen, milking parlor line-ups, all of which could potentially contribute to the increase in incidence rate (Sage et al., 2024; EFSA et al., 2025).

In this study, we present a validated stochastic agent-based model to estimate the epidemiological parameters for intra-herd HPAI transmission in Californian Dairy herds. The hierarchical agent-based model also parameterized stochastic cattle movements within-herd to simulate real dairy management practices. This heterogenous transmission model was optimized using observed HPAI incidence on four commercial dairy farms from the Central Valley of California, USA.

## METHODS

### Hierarchical agent-based simulation model

To simulate the outbreak of HPAI, we developed a hierarchical agent-based simulation model. The detailed explanation of the model is provided in the Supplementary Material: Overview design details (ODD) protocol, for reproducibility in accordance with the guidelines for describing agent-based models (Grimm et al., 2020). All codes and protocols were developed in Python 3.11 using object-oriented programming to generate agentic hierarchy (Miller et al., 2015). Here, we provide an overview of the model.

In this model, the fundamental agents were cows (Level 1), followed by the pens that housed these cows (Level 2/ Aggregate) and finally the whole farm (Level 3/ Aggregate). Each agent from each tier had specific attributes and sub-functions that contributed to the simulation.

We sourced data for within-farm cattle movements between different pens from four mixed breed (Holstein and Jersey) Central Valley farms. This secondary data was used in a previous study (Konboon et al., 2018) to map the dwelling time ranges of cows in each of the functional pens on the dairy farm. We used these dwelling time ranges as inputs for our model, along with other parameter inputs seen in Supplementary Tables S4 and S5. Our model simulated intra-farm movement of cows between different functional pens by adapting a modified version of the pen movement framework (Konboon et al., 2018). Our adaptation showed an introduction of a hospital pen which is standard practice in most dairy herds of California’s Central Valley. Additionally, we masked the calf and heifer pens since these animal units were housed on external sites. The adapted pen movement framework can be seen in Figure 1. For each cow, the dwell times for the pens were stochastically sampled from the minimum and maximum duration (Figure 1)

**Figure 1.**
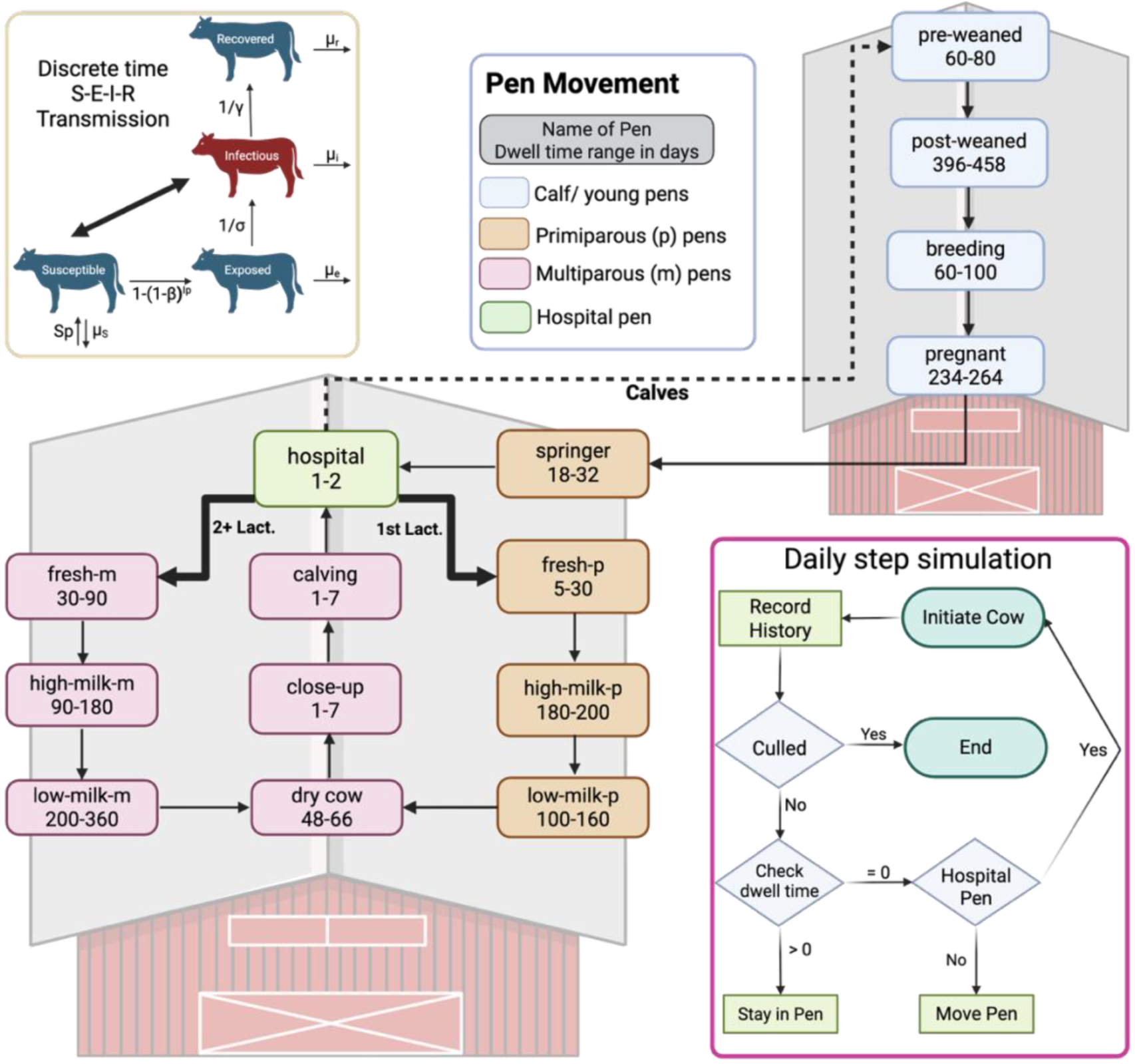
Schematic diagram of Hierarchical Agent-Based Model used for simulating the intra-farm outbreak of HPAI in Californian Dairy Herds.

Along with the movement module, an epidemiological S-E-I-R (Susceptible-Exposed-Infectious-Recovered) module was integrated. In every time step (daily), the cows housed in the same pens interacted with each other. Based on the epidemiological parameters such as transmission rate (β), recovery rate (γ), incubation period (T_E_) and number of infectious (I_p_) and susceptible cows (S_p_) in any pen, the cows transmitted HPAI infection wherein the susceptible cows became “exposed”. The probability of exposure, P(exposure), was based on discrete time sampling giving the number of exposed cows E_p_. Each cow also had a probability of being culled (*P*(*cull*)*^t^*) in each time step giving μ_annual_, the annual culling rate of the dairy herd (Eq. 1). In each pen, at every time step, the number of cows in a particular epidemiological state (S, E, I, R) were recorded using the discrete time transmission model formulation as shown below (Eq. 2–5)

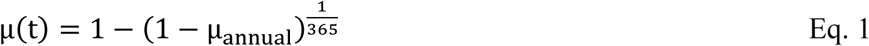

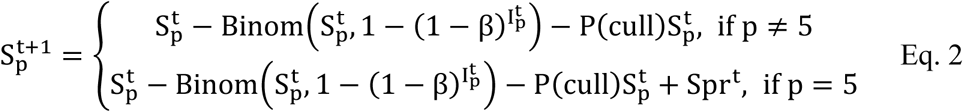

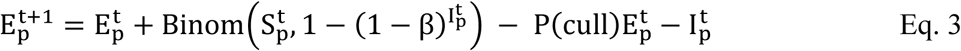

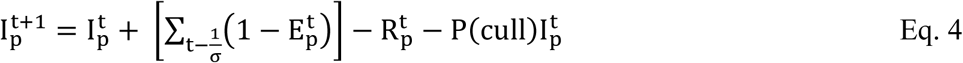

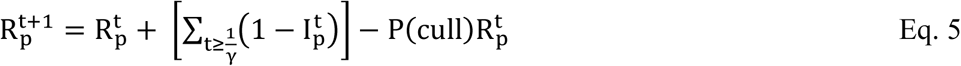

Where 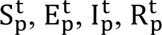 are number of susceptible, exposed, infectious and recovered cows in pen p at times t respectively. β is the transmission rate, 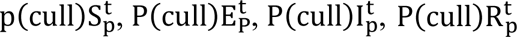 are the proportion of culled susceptible, exposed, infectious and recovered cows at time t. New susceptible cows/ springers (Spr^t^) are also added when they enter the Springer pen (Figure 1). 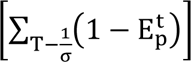 are the proportion of exposed cows that turn infectious at time t at the end of their incubation period 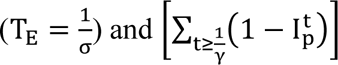 are all the cows that are recovered by the time t with recovery rate of γ.

On the herd level, the total number of cows in each epidemiological state is summed up from each pen (Eq. 6)

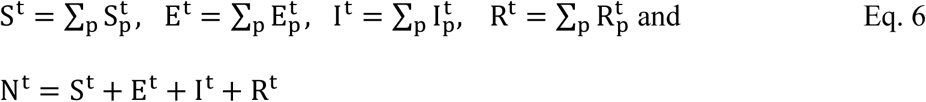

Where N^t^, S^t^, E^t^, I^t^, R^t^ are the herd size, proportion of susceptible, exposed, infectious and recovered cows in the herd respectively at time t, and pen p ɛ {Springer…Hospital}.

### Bayesian Optimization

We sourced case data on four Central Valley dairy farms between September and October of 2024 from an ongoing longitudinal study conducted by the Veterinary Medicine Teaching and Research Center (VMTRC, University of California Davis). All four of these farms had reported lactating cows that presented clinical signs during this time. All four farms had calf rearing sites outside of the main facility (open farms). The calves were reared at the rearing site, and springers close to calving were introduced to the main herd. One of these dairies raises their heifers on site from approximately four months of age. The herd sizes of two farms were small (∼500 and ∼800 lactating cows respectively). The remaining two farms had larger herds (∼2500 lactating cows each). In order to simulate the intra-farm HPAI outbreak as close to reality as possible using the reference case data, we fitted a Bayesian Optimization (BO) algorithm that used a surrogate model with Gaussian Processes (BO–GP) with expected improvement acquisition that minimized the difference between simulated cases and target cases that were sampled from the farms. Since the target cases were collected only from the milking cows, our simulated case data was filtered to include counts of the cows that turned infectious in milking pens (High Milk primiparous, Low Milk primiparous, High Milk multiparous and Low Milk multiparous). BO–GP is an efficient optimization protocol that uses surrogate models (Gaussian models) to approximate the objective function (mean and uncertainty) while minimizing the difference in each iteration (Shahriari et al., 2016). While BO is normally applied to hyperparameter tuning of deep learning models instead of operational optimization, it was found to be flexible to be suited for our purpose (Blanks and Brown, 2024).

### Loss Function

Considering that the case data collected could be construed as a time series inside the total duration of the outbreak as well as count data, we developed two different loss functions for optimization. Firstly, a Distance Metric (DM) that combined Dynamic Time Warping (DTW), Mean Absolute Error (MAE) and Cross-Correlation based time lag error (CC) was used to optimize the simulated case time series against the target case time series. This distance-based metric has been previously reported to efficiently match time series and sequential count data operations (Lee et al., 2024). Secondly, we developed a Poisson Loss Function (PLF) that assumes Poisson distribution for target case data at each time point. Using penalized quasi-likelihood maximization, the PLF are well-suited for case counts (Vazquez et al., 2009).

### Protocol and objective function

Four parameters, viz., Transmission rate, Recovery rate, Incubation Period, and number of initially introduced infectious cows were optimized using the BO–GP setup. The optimization protocol was integrated in the hierarchical agent-based simulation model. For each iteration, parameter values were sampled from a bounded hyperspace for the four parameters at random and the loss function (either DM or PLF) was calculated. BO–GP was parameterized with 1000 iterations with a convergence error of 0.0001 and 50 random starts to avoid local optima. The algorithm converged when minimum possible loss was reached and no improvement (decrease in difference between simulated and target data) was possible.

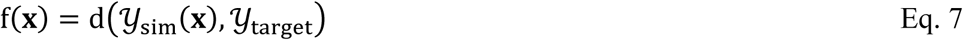

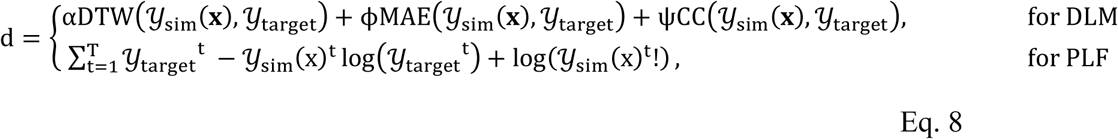

Where, *α*, *φ*, *ψ* are the weighting coefficients of DTW, MAE and CC time lag respectively. Y_sim_(**x**) is the simulated case data using **x** parameter values for the four parameters and Y_target_ is the target case data. arg max

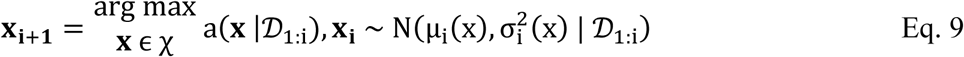

Where χ is the bounded hyperspace of possible values for the set of parameters (**x**), D_1:I_ is the combination of sampled parameter values and the loss metric calculated between simulated data and target data until iteration i (updated prior), a(x |D_1:i_) is an acquisition function that represents the expected decrease in the loss using the surrogate model (GP) for **x_i_**_+**1**_ based on the updated prior performance of D_1:I_ for the newly predicted values in **x_i_**_+**1**_with an estimated mean (μ_i_(x)) and variance 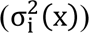 by the surrogate model (GP) based on D_1:i_.

We assumed that the true functional association between the parameter values and the loss metric between target and the simulated cases was unknown (Eq. 7). Depending on the loss function used (Eq. 8), the objective function aimed to mimic the HPAI outbreak using the target case data in the simulation model (Eq 9). The algorithm and the custom loss functions and the wrapper for the objective function were all coded in Python using gp_minimize() function from Scikit-Optimize 0.8.1 package (Holger Nahrstaedt, Gilles Louppe, Manoj Kumar).

### Effective transmission rate and basic reproduction number

By simulating the optimal values of the four parameters (N_iters_ = 100 for each farm), we calculated the effective transmission rate using Eq. 10. To estimate the overall effective transmission rate, we averaged (mean) the transmission rate per day of the observed outbreak.

Using this effective transmission rate, we derived the basic reproduction number (R_0_) for intra-farm transmission (Eq. 11)

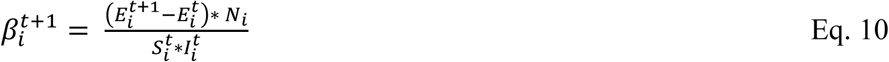

Where 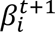 is the effective transmission rate on day 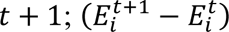 is the incidence on that day, *N_i_* is the herd size and 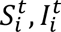 are the number of susceptible and infectious cows on farm *i*.

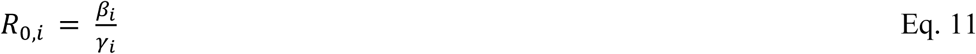

Where *R*_0,*i*_, *β_i_*, *γ_i_* are the basic reproduction number, average daily effective transmission rate, recovery rate respectively on farm *i*.

### Sensitivity Analysis

We performed sensitivity analysis on the optimized parameters of transmission rate, recovery rate, incubation period and initially introduced infectious cows. The target outcomes were peak infectious cows count, cumulative infectious count, cumulative exposed count, final count of recovered (unculled) cows. Additionally, we also tested the sensitivity of the four outcome variables to the changes in herd size of the simulated farm. We performed both global and local sensitivity analysis in base Python 3.11 and with the help of SALib package (Iwanaga et al., 2022) which is specifically used for sensitivity analysis.

### Global Sensitivity analysis

We performed Morris’ sensitivity analysis (Morris, 1991) for initial exploration of the importance of the five (epidemiological + herd size) parameters on the four outcome variables. Morris’ method, being computationally less expensive than the variance-based method was suitable for large parameter value hyperspace. The Morris’ method was parameterized with 20 trajectories (r = 20; total runs = r × (k + 1) = 20 × 6 = 120, where k is the number of parameters). The mean effect and absolute effect along with the standard deviation were estimated. Finally, we performed Sobol analysis (Sobol, 2001) on the five parameters to evaluate their contribution (both first order and interactions) on the variance of the four outcome variables. This analysis was performed to provide rigorous information on the importance of the parameter values, especially considering that an interaction effect was highly likely (Edwardes et al., 2024). We generated sample sets for Sobol analysis using Saltelli quasi-random sampling algorithm (Saltelli, 2008; Saltelli et al., 2010) with a sample size of 256 samples per run (N = 256, total runs = N × (2k + 2) = 3072, where k = 5 parameters). The first order (main), second order (pairwise interaction) and the total order effects were evaluated for the four outcome variables.

### Local Sensitivity analysis

We performed local sensitivity analysis on all possible values of the five parameters to estimate the differences in the four outcome variables when compared to the optimal values of the parameters drawn from the BO (optimal value of herd size was fixed at 500). We used One-At-a-Time (OAT) method to vary the values of one parameter at a time while holding others constant to test the local sensitivity.

## RESULTS

### Estimated Epidemiological Parameters

We estimated basic reproduction numbers (R_0_), calculated using SEIR simulations done for four observed Californian herds for which transmission parameters were estimated using a novel BA-GP method (Figure 2A). Overall, the estimated R_0_ suggested high variation in the transmission intensity across different farms. Average R_0_ on all farms based on the first 15 days of outbreak were relatively high, indicating intense transmission early in the outbreak. Farm3 consistently had the highest mean R_0_ throughout the observed outbreak and with both estimation methods, suggesting the most transmissible farm setting. Farm 2 R_0_ was generally the lowest, implying farm-level differences in initially introduced infectious cows, contact patterns or other mechanisms.

**Figure 2.**
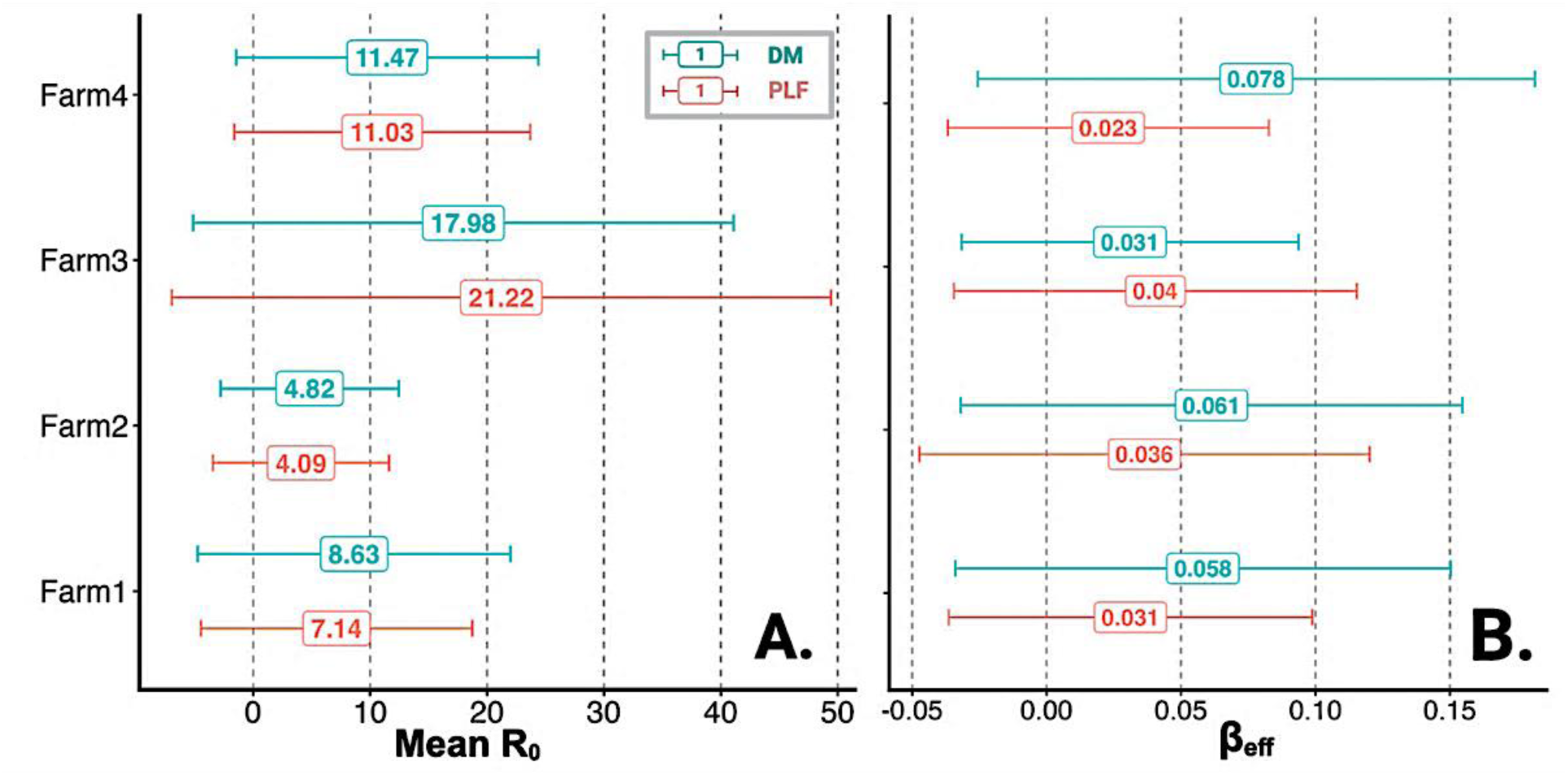
(A) Estimated basic reproduction number (R_0_) based on first 15 days in outbreak and (B) the estimated effective transmission rate *β_eff_*.

Overall, the estimated average R_0_ was in the range of 10.7-10.8 across all farms and outbreak time frames with substantial variation around the mean (± 4 SE) depending on the optimizer function (averaged across all farms in Figure 2A). Farm 3 shows the highest mean R_0_ (DM: 18, PLF: 21.2) and the greatest variability and Farm 2 consistently had the lowest mean R_0_ (DM: 4.8 PLF: 4.1). Estimated R_0_ values based on Poisson Loss Function (PLF) optimization were similar to or slightly lower than the Distance Metric (DM) based optimizer early in the outbreak (1-15 days of outbreak), signaling that method choice strongly influenced the estimated epidemiological parameters.

The overall effective transmission rate calculated using Eq. 10 was 0.045 (± 0.008) across all four farms (Figure 2B). The simulated SEIR curves using the average daily R_0_ from the two optimizers, showed two peaks for exposed and infectious cows signaling bimodal transmission and re-introduction of infectious cows in the herd (Figure 3; Susceptible counts not shown).

**Figure 3.**
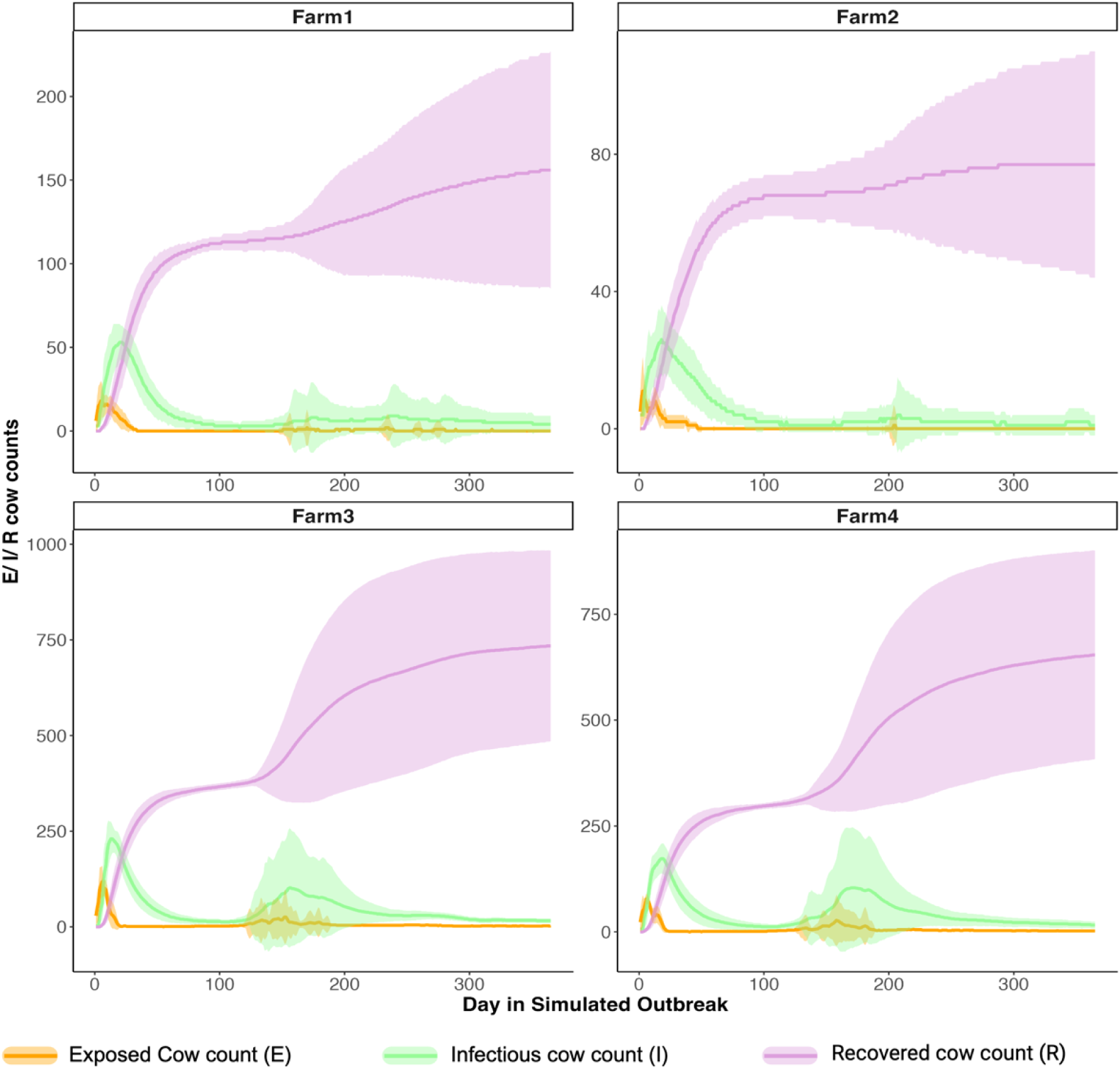
Simulated SEIR counts for each of the modeled farms over one year of outbreak.

### Bayesian Optimization using Gaussian Process (BO–GP)

The posterior distribution prior to the optimization using BO–GP narrowed the uniform priors for the four parameters. The posteriors represented the top 5% of simulated outbreak curves for each of the modeled farms based on the DM and the PLF metrics (see Supplementary Figure S1). These posterior ranges were between 0.01 to 0.1 successful contacts per cow per day for transmission rate parameter, 0.04 to 0.08 cows per day for recovery rate parameter, 1 to 10 days for incubation period and 1 to 10 cows for initially introduced infectious cows for the four parameters. The hyperspace of values for these ranges was used for optimization. BO–GP optimization using DM and PLF yielded slightly different optimal values for the four parameters for the outbreak on each farm.

The optimal values remained consistent for the transmission and recovery rates and differed slightly for incubation period and initially introduced infectious cows based on which metric was used in optimization (Figure 4 and Figure 5C). Post-optimization simulations (N = 100 for each farm) showed that the optimized values of parameters generated a simulated curve that was moderately similar to the real incidence data (target outbreak) on the four farms. The simulated outbreak curves were better fits for Farms 3 and 4 but not for Farms 1 and 2 (Figure 4 and Figure 5C). The optimal values based on PLF followed the trend of the incidence data slightly better than the DM based optimal values. On average, for DM-based the internal transmission and recovery rates across all four farms were estimated at 0.015, 0.055 respectively. The average for DM-based optimal values of incubation period and initially infectious cows were estimated at 2.5 days and 3 cows respectively. The PLF-based optimal values were estimated to be 0.01, 0.055, 2.25 days and 8 cows respectively.

**Figure 4.**
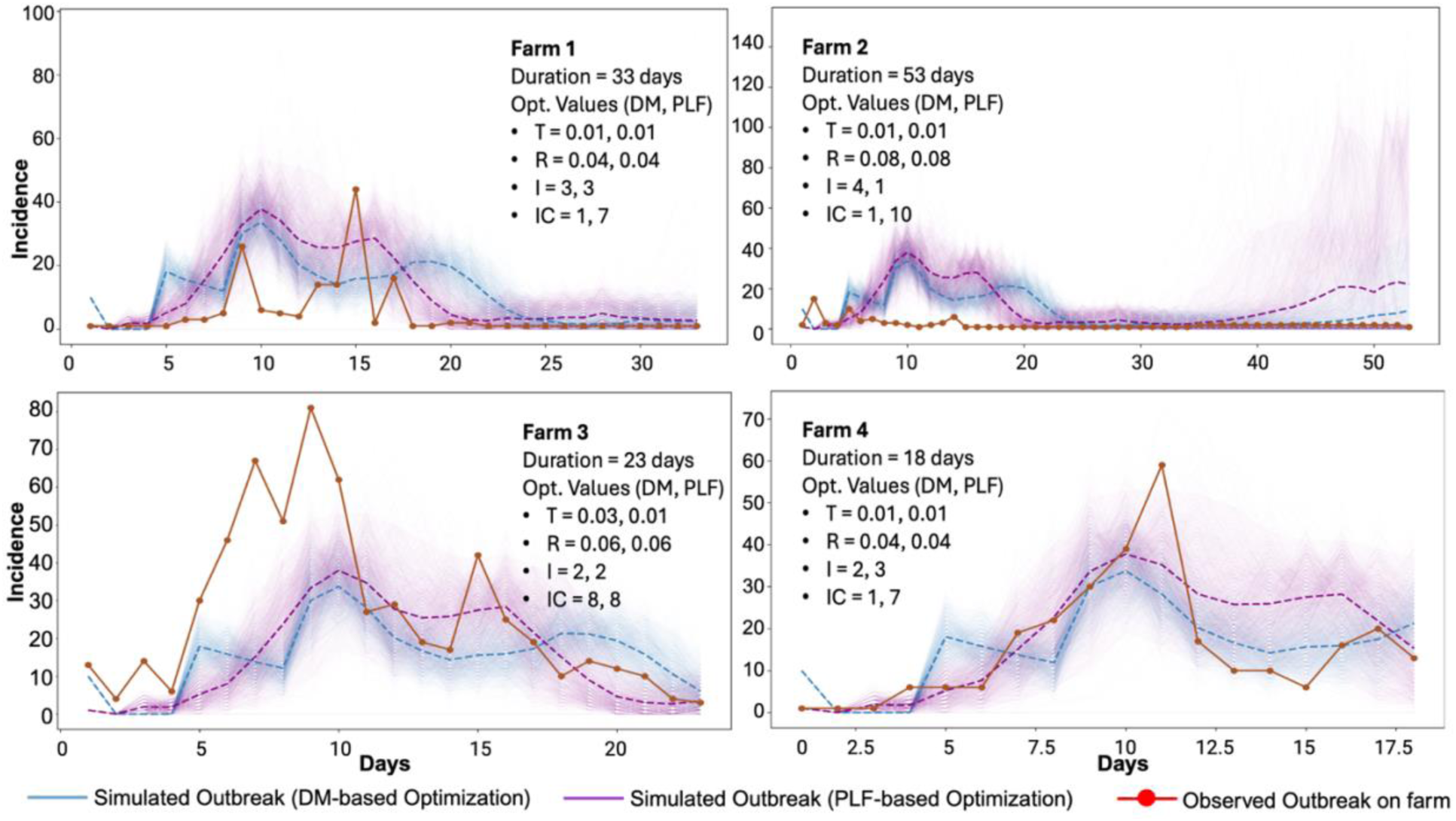
Optimized vs. Observed outbreak incidence curves for four modeled farms using DM (Distance-based Metric) and PLF (Poisson Loss Function) metrics for optimization using BO–GP algorithm. *Abbrev. In figure: T = Transmission Rate Parameter, R = Recovery rate, I = Incubation Period in days, IC = Initially introduced infectious cows

**Figure 5.**
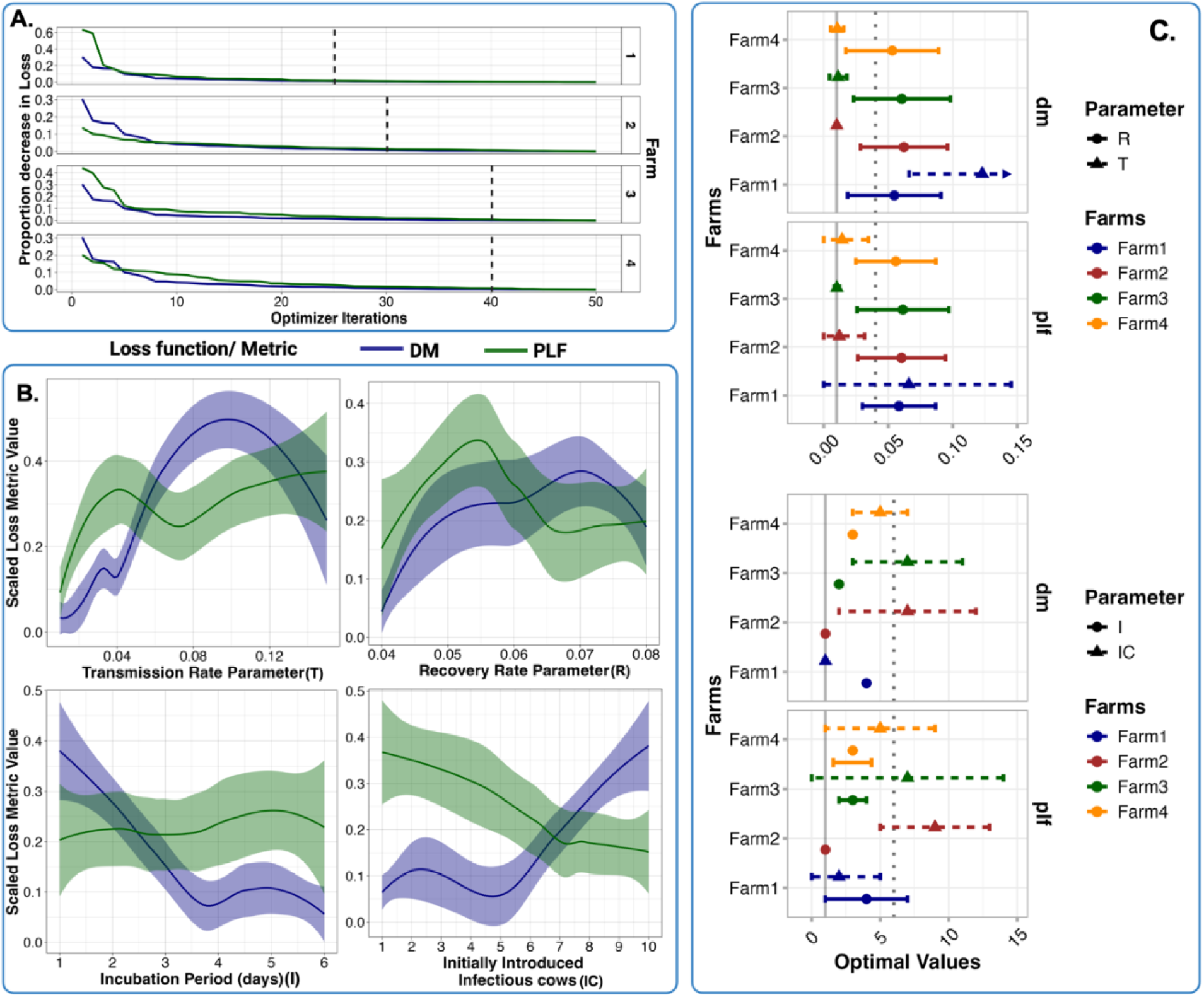
Optimization results illustrating (A) Convergence using two loss function metrics, (B) relationship of parameter values with loss function metrics across all four farms and (C) Averages of optimized parameters with 95% confidence intervals.

From Figure 4 and Figure 5C, for Farm 1, the optimized values for transmission and recovery rates, incubation period and number of initially introduced infectious cows were found to be 0.01, 0.04, 3 days and 1 cow for DM-based optimization respectively. The PLF-based optimization yielded the same optimal values except for number of initially introduced infectious cows (7 instead of 1). For Farm 2, the optimized values were 0.01, 0.08, 4 days and 1 cow respectively from DM-based optimizer. The PLF yielded a difference in value for incubation period and initially infectious cows with values of 1 day and 10 cows respectively. For Farm 3, the DM-based optimal values were 0.03, 0.06, 2 days and 8 cows respectively. PLF-based values differed from DM-based values for transmission and recovery rates (0.01 and 0.06 respectively). For Farm 4, DM-based values were 0.01, 0.04, 2 days and 1 cow. PLF-based values for incubation period and initially infectious cows were 3 days and 7 cows respectively that differed from DM-based values.

BO–GP optimizer converged for farms 1, 2, 3, and 4 within 25, 31, 41 and 41 iterations respectively (Figure 5A). No specific association between the values of the four parameters and either of the loss function metrics were found across the four farms (Figure 5B). For transmission rate, the lower values within the posterior range were associated with higher loss metric for PLF-based optimizer compared to DM-based optimizer. This pattern was similar for recovery rate.

For incubation period, the loss consistently decreased with increase in parameter value for DM-based optimizer but remained constant for PLF-based optimizer. For number of initially infectious cows, the loss associated with parameter values showed opposite trends (increase vs decrease against increase in number of cows) for DM-based and PLF-based optimizers

### Sensitivity analyses

We tested the sensitivity of four outcomes, namely, (i) peak infectious count, (ii) total/ cumulative infectious cow count, (iii) total/ cumulative exposed cow count (where ii+iii is the cumulative incidence), and (iv) final recovered cow count against the value range for four optimized parameters (transmission and recovery rates, incubation period, number of initially introduced infectious cows) with an additional parameter of herd size of the farm on local and global levels. Local sensitivity results showed that all four outcomes were sensitive to parameters such as number of initially introduced infectious cows, herd size and recovery rate of cows (Figure 6A).

**Figure 6.**
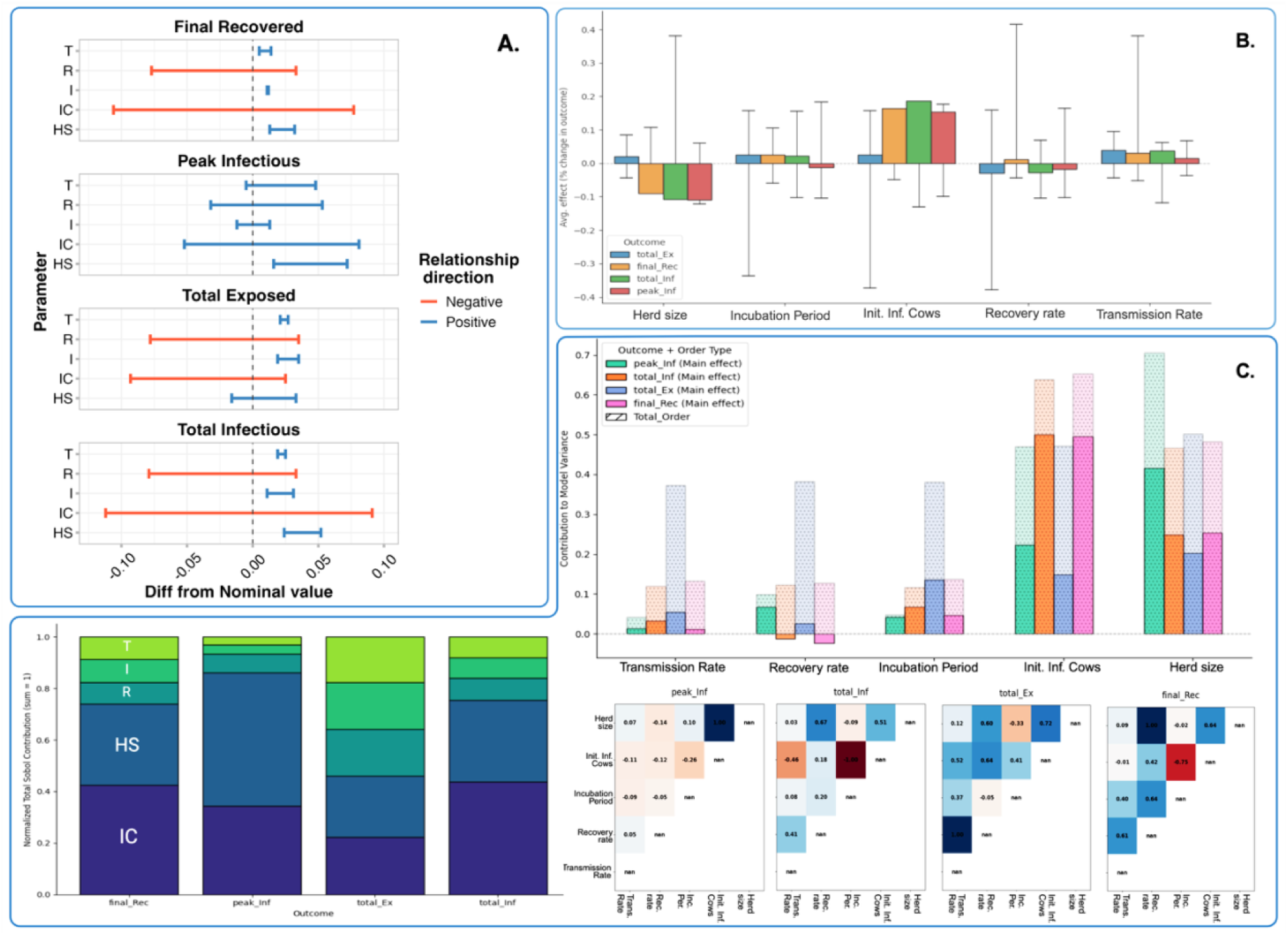
Sensitivity analyses outcomes for peak infectious count, total infectious count, total exposed count and final number of recovered cows against the values of optimized four parameters and herd size on (A) local scale and (B) global sensitivity analyses using Morris’s method and (C) Sobol method showing total and second order (interaction) sensitivity. *Abbrev. in figure: T = Transmission Rate Parameter, R = Recovery rate Parameter, I = Incubation Period in days, IC or Init. Inf. Cows = Initially introduced infectious cows, H = herd size of the farm, peak_Inf = peak infectious cow count, total_Inf = total/ cumulative infectious cow count, total_Ex = total/ cumulative exposed cow count, final_Rec = final recovered cow count

Outcomes were found to be locally robust to changes in transmission rate and incubation period showing that the majority of sensitivity stemmed from the number of cows that remained infectious rather than rate of transfer of infection between interacting cows. Global sensitivity analysis showed similar results wherein all four outcomes were found to be sensitive to the number of initially introduced infectious cows, herd size and recovery rate. The variance in outcomes was found to be robust against changing values of transmission rate and incubation period (Figure 6B and 6C).

## DISCUSSION

The aim of this study was to estimate the outbreak parameters of HPAI in dairy herds. Our model framework simulated the HPAI outbreak in four California dairy herds by calibrating the model parameters so as to mimic the incidence data. Our modeling framework replicated the within-herd dynamics such as movement of cows within different pens, lactations, culling and replacement as well as HPAI cattle-to-cattle transmission of the four farms under study. Unlike conventional models, this model is independent of mechanistic or probability distribution assumptions. Due to built-in fast optimization and high accuracy, varied scenarios are easily testable. The model is easily integrated in decision-making for vaccination and cost-effectiveness of disease monitoring.

We estimated that 4.5 per 100 daily contacts between susceptible and infectious cows within the lactating pens resulted in a successful transmission of infection. Our overall estimated R_0_ of 10.75 across different timeframes over the first 15 days of the observed outbreak within all four modeled farms was moderately higher than the estimate of 8 (range of 6-10 over different scenarios) by (Malladi et al., 2026) signaling a fundamental difference in the data from various sources.

Our model demonstrated that the movement of cows between pens ensured a localized cluster of outbreaks within the sub-herds (pen population) that prolonged the overall outbreak within farms. We estimated the total duration of infection (incubation period + duration of infectiousness) between 14.5 and 28 days (estimated range averaged over four farms with two optimization metrics), which is higher than the estimates from the homogenous models of (Bellotti et al., 2024) but within the margins of the overall field estimates of 10-45 days for duration of the epidemic in affected US herds (Rodriguez et al., 2025; Malladi et al., 2026).

In highly heterogenous systems such as disease transmission between herd-mates along with pen movements, multiple parameter combinations can generate similar outcomes (Norton et al., 2025). Owing to this uncertainty, an optimal “best-fit” solution is difficult to estimate using descriptive agent-based models. Reproduction of observed outbreaks by agent-based models through a set of equifinal parameter combinations cannot be purported to be the same as reproduction of the actual system dynamics which is why appropriate model calibration is necessary (Box, 1979). Using BO–GP which is a surrogate approach to calibration of the simulated outbreak, our framework iteratively samples parameter combinations that are likely to match the observed outbreak thereby optimizing parameters without assuming any mechanistic underpinnings. Notably, our model yielded similar outbreak curves, especially for farms with substantial incidence (Farm 3 and 4) by narrowing the uncertainty within parameters. For Farms 1 and 2 where lower incidence was reported, our model overestimated the number of infectious cows which is consistent with the behavior of similar models due to poor data quality or possibly uncaptured heterogeneity (Kirkeby et al., 2021; Leung et al., 2023). To date and to our knowledge, only a few studies have attempted to use GP based Bayesian inference in disease transmission modeling and only one study on Malaria transmission has integrated hierarchical agent based modeling with BO–GP optimization (Reiker et al., 2021).

Considering that the cows from the same pen are milked together (Guinan et al., 2025) and that preliminary reports pointed to HPAI transmission primarily occurring at the site of milking (Burrough et al., 2024; Kamel et al., 2025), it is possible that the lactating cows’ pens would become hotspots for HPAI outbreak within the herd. However, considering that we only counted the incidence in lactating cows, the effective transmission rate could have been underestimated when not considering the dynamics in pens such as calving and hospital pens. The modeling choice to focus on lactating cows was in line with the target incidence data that was collected based on reporting of clinical signs in cows by farm workers during milking (Sharif S. Aly; personal communication). This was consistent with nationwide testing for seropositive samples that sampled lactating cows (Caserta et al., 2024; Lombard et al., 2025). Since the sampled data was based on preliminary incidence reports that farm workers made based on cows presenting with clinical signs during or before milking, it was prudent to assume that the asymptomatic cows were not included in the target case curves which our models aimed to emulate.

We fit an individual level SEIR model (Susceptible-Exposed-Infectious-Recovered) which differed from the compartment-based SIR model (Susceptible-Infectious-Recovered) used in other studies (Bellotti et al., 2024). We intended to capture the latency between when cows are exposed and when they start showing clinical signs that can be reported by farm workers (Rodriguez et al., 2024) in the absence of active surveillance data. Due to the pen-level movements of cows and heterogeneity in interaction between herd mates, we chose to calibrate hierarchical agent-based models for simulating intra-farm dynamics of HPAI outbreaks over homogenous compartmental models. Since the nationwide HPAI dairy outbreak was an emergent occurrence without long-term prevalence data, agent-based modeling architecture facilitated the capture of population level disease spread and improvement in the understanding of the contributing factors by validating against preliminary incidence data based on clinical signs (Hunter et al., 2017).

The sensitivity of our model to initially introduced infectious cows, recovery rate and herd size was consistent with similar SEIR modeling frameworks for infectious diseases (Skrip and Townsend, 2019; Muhammad et al., 2021; Ma et al., 2022). Further, from sensitivity analysis, our models were robust to changes in transmission rate and incubation period on both local and global scales. This demonstrated that varied structure of within-herd movement, pen structure or increased contact between herd-mates would not, in theory, affect the outbreak size (peak incidence). Higher sensitivity of incidence to herd size and initial introduction of infectious cows (Figure 6) meant that larger herd sizes paired with open herd operations and a high level of cattle movement would increase the risk of bigger outbreaks. Moderately high sensitivity of outcomes to recovery rate (Figure 6) signaled that effective intervention and timely treatment would possibly reduce the size and duration of the outbreak. However, these sensitivities were not tested for specific conditions such as localized superspreaders within pens or quarantine of clinically symptomatic cows which would ensure changes in contact rates that would alter the simulations (Nsoesie et al., 2012).

Additionally, our modeling framework is amenable for testing different mitigation strategies and future prevention effectively and on a granular scale by targeting hotspots of outbreak within modeled farms (Skrip and Townsend, 2019). Multi-objective decision frameworks such as vaccination and monitoring compliance on individual cow-level, pen-level and herd-level along with economic and social incentives can be feasibly tested by adapting such modeling frameworks (Sok and Fischer, 2020).

## CONCLUSION

The integration of Bayesian optimization, and intra-farm simulation using a hierarchical agent-based model combined with robust sensitivity analysis, produced actionable insights on intra-farm spread of HPAI within cow herds, guiding both future model development and applied disease control strategy.

## NOTES

### Credit authorship contribution statement

P.S.K: Conceptualization, methodology, software, formal analysis, writing (original draft); S.S.A: validation, resources, writing (review and editing), supervision; D.R.W: Resources, investigation, writing (review and editing); W.R.E: Resources, investigation, writing (review and editing); P.S.P: Conceptualization, methodology, funding acquisition, writing (original draft), writing (review and editing), project administration, supervision.

### Data availability statement

Access to raw animal-related and disease-incidence data is restricted to protect confidentiality and to comply with the terms of collaborative agreements and institutional permissions. Requests regarding potential access to the raw data should be directed to the UC Davis Veterinary Medical Teaching and Research Center (VMTRC). All intermediate processed datasets, analytical workflows, model code, and the supplementary materials necessary to reproduce the analyses in this manuscript are publicly available on Github: https://github.com/EpiPandit/Kulkarni_et_al_HPAI_Dairy_Cows/tree/main

### Ethics statement

All animal-related and disease-incidence data used in this study were anonymized secondary data collected by the UC Davis Veterinary Medical Teaching and Research Center (VMTRC) as part of routine clinical recording and operational activities, under existing collaborative agreements and permissions. No human participants or live animals were used for this analysis so, this study did not require review or approval by an Institutional Animal Care and Use Committee or Institutional Review Board.

## Supporting information

Supplementary Results

Supplementary Files ODD Protocol

