## Supplementary Results for "Estimation of the transmission dynamics of H5N1 HPAI outbreak in a dairy herd using a modeling approach"

### Supplementary material: Additional Results

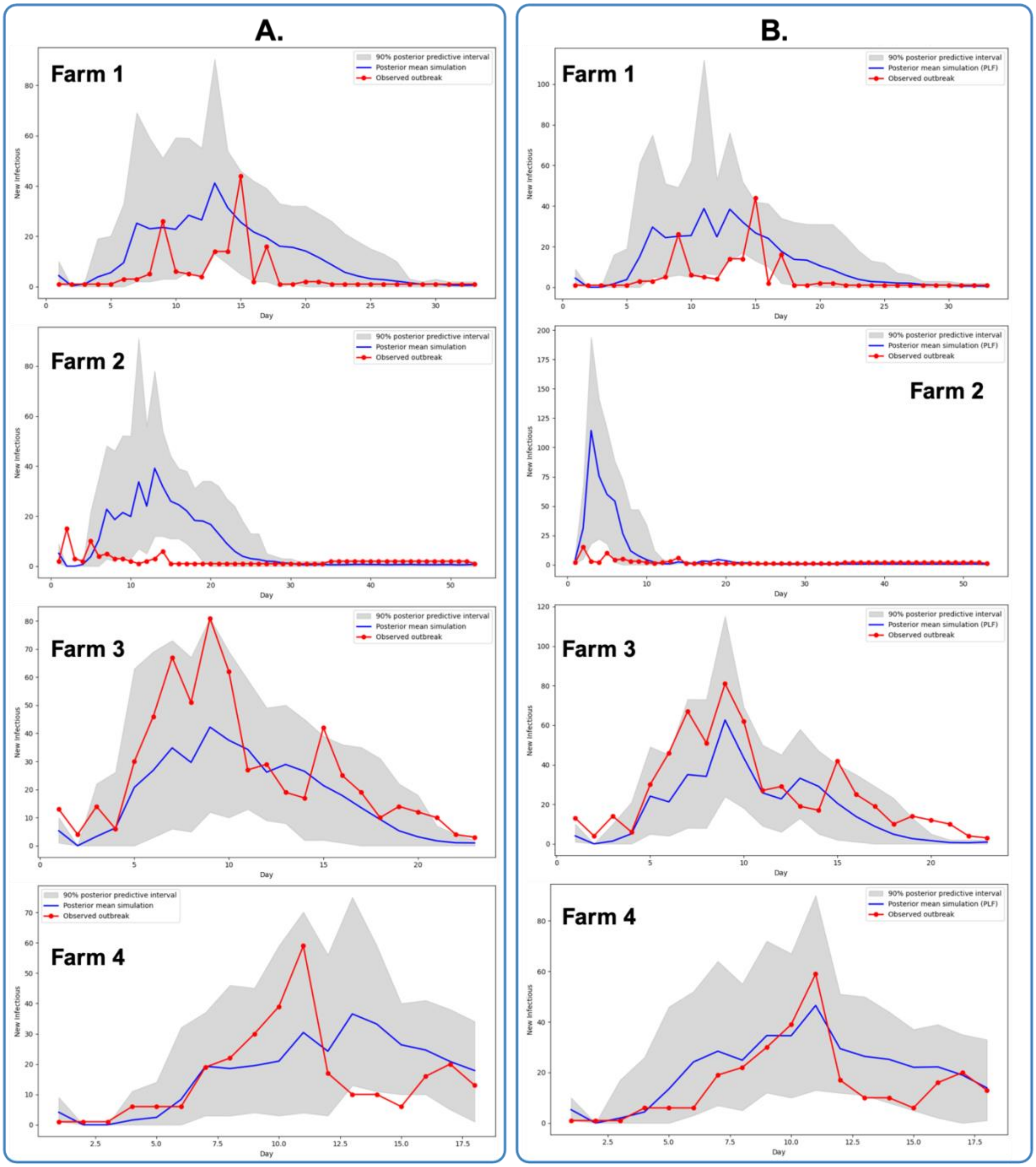

### Estimation of the transmission dynamics of H5N1 HPAI outbreak in a dairy herd using a modeling approach

**Pranav S. Kulkarni , Sharif S. Aly , Deniece R. Williams , Wagdy R. ElAshmawy ,**

**Pranav S. Pandit**

**Figure S1. Simulation results (N = 200 per farm) of intra-farm outbreak using posterior ranges for transmission and recovery rates, incubation period and number of initially introduced infectious cows using (A) Distance-based metric (DM) and (B) Poisson Loss Function (PLF)-based metric.**

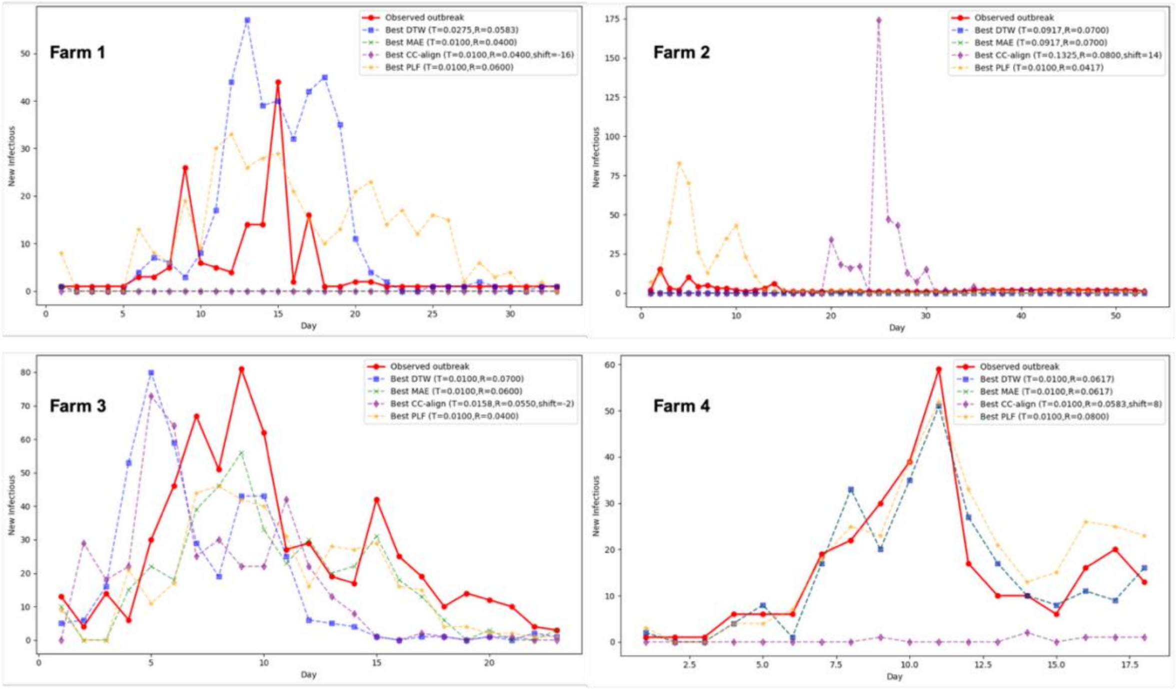


**Figure S2. Post-optimization simulation of outbreak against observed outbreak on 4 modeled farms for each separate Distance-based Metric (DM) and Poisson Loss Function based metric used in optimization. Abbrev. in figure: DTW= Dynamic Time Warping, MAE= Mean Absolute Error, CC_align= Cross Correlation temporal Alignment/ shift, PLF= Poisson Loss Function, T = Transmission rate, R = Recovery rate.**

### Estimation of the transmission dynamics of H5N1 HPAI outbreak in a dairy herd using a modeling approach

**Pranav S. Kulkarni , Sharif S. Aly , Deniece R. Williams , Wagdy R. ElAshmawy , Pranav S. Pandit**

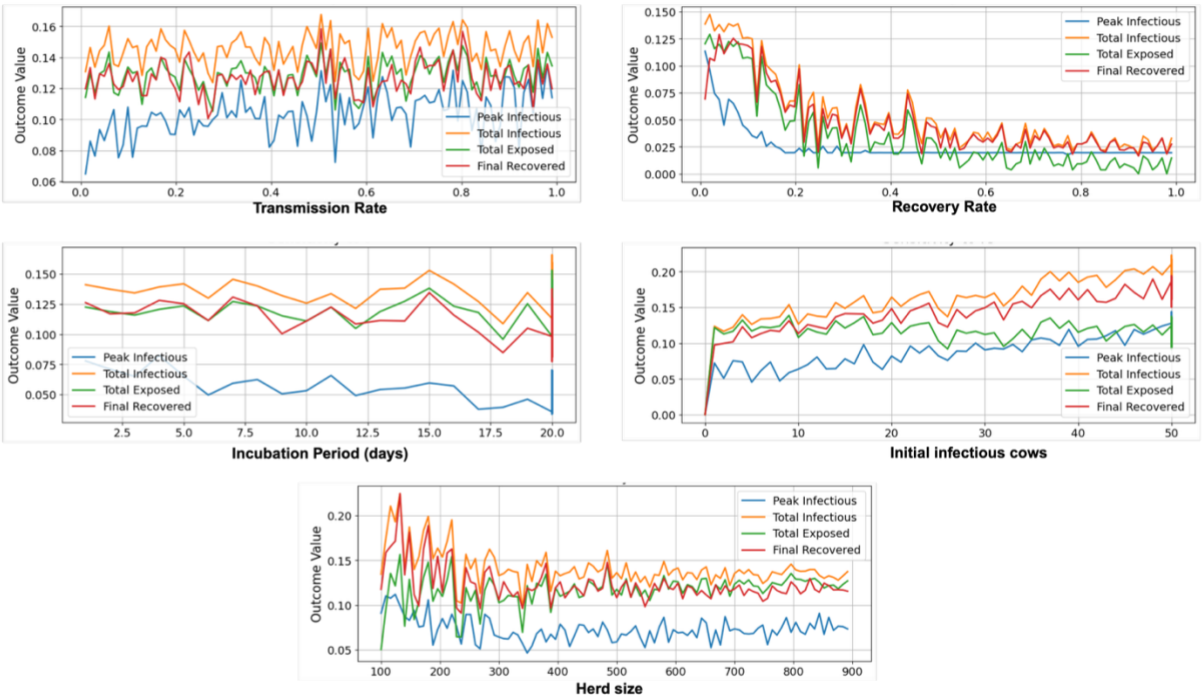


**Figure S3. One-way Local Sensitivity results of four outcomes (Peak and Total/ cumulative Infectious counts, Total Exposed count, Final Recovered count) against full range of values for four parameters (transmission and recovery rates, incubation period and number of initially introduced infectious cows) and additional input of herd size.**

**Table S6. Descriptive statistics of Morris Sensitivity analyses with 95% confidence intervals for 4 outcomes against 5 input variables**

| **Outcome (Cow Counts)** | **Absolute Average Effect** | **Std. Dev.** | **Average Effect** | **Lower 95% CI** | **Higher 95% CI** |
|---|---|---|---|---|---|
| **Herd size** | | | | | |
| Total Exposed | 0.03 | 0.03 | 0.02 | -0.04 | 0.09 |
| Final Recovered | 0.09 | 0.12 | -0.09 | -0.34 | 0.16 |
| Total Infectious | 0.11 | 0.13 | -0.11 | -0.37 | 0.16 |
| Peak Infectious | 0.11 | 0.13 | -0.11 | -0.38 | 0.16 |
| **Incubation Period** | | | | | |
| Final Recovered | 0.03 | 0.03 | 0.03 | -0.04 | 0.10 |
| Total Exposed | 0.03 | 0.04 | 0.03 | -0.06 | 0.11 |
| Total Infectious | 0.03 | 0.04 | 0.02 | -0.06 | 0.10 |
| Peak Infectious | 0.01 | 0.02 | -0.01 | -0.05 | 0.02 |
| **Number of initially introduced Infectious cows** | | | | | |

### Estimation of the transmission dynamics of H5N1 HPAI outbreak in a dairy herd using a modeling approach

**Pranav S. Kulkarni , Sharif S. Aly , Deniece R. Williams , Wagdy R. ElAshmawy , Pranav S. Pandit**

| | | | | | |
|---|---|---|---|---|---|
| Total Infectious | 0.19 | 0.11 | 0.19 | -0.04 | 0.42 |
| Final Recovered | 0.16 | 0.11 | 0.16 | -0.05 | 0.38 |
| Peak Infectious | 0.15 | 0.11 | 0.15 | -0.07 | 0.38 |
| Total Exposed | 0.05 | 0.06 | 0.03 | -0.10 | 0.15 |
| **Recovery rate** | | | | | |
| Final Recovered | 0.04 | 0.07 | 0.01 | -0.13 | 0.15 |
| Peak Infectious | 0.02 | 0.04 | -0.02 | -0.11 | 0.07 |
| Total Infectious | 0.04 | 0.04 | -0.03 | -0.12 | 0.06 |
| Total Exposed | 0.04 | 0.05 | -0.03 | -0.12 | 0.06 |
| **Transmission Rate** | | | | | |
| Total Exposed | 0.06 | 0.07 | 0.04 | -0.10 | 0.18 |
| Total Infectious | 0.05 | 0.07 | 0.04 | -0.10 | 0.18 |
| Final Recovered | 0.05 | 0.07 | 0.03 | -0.10 | 0.16 |
| Peak Infectious | 0.02 | 0.03 | 0.01 | -0.04 | 0.07 |

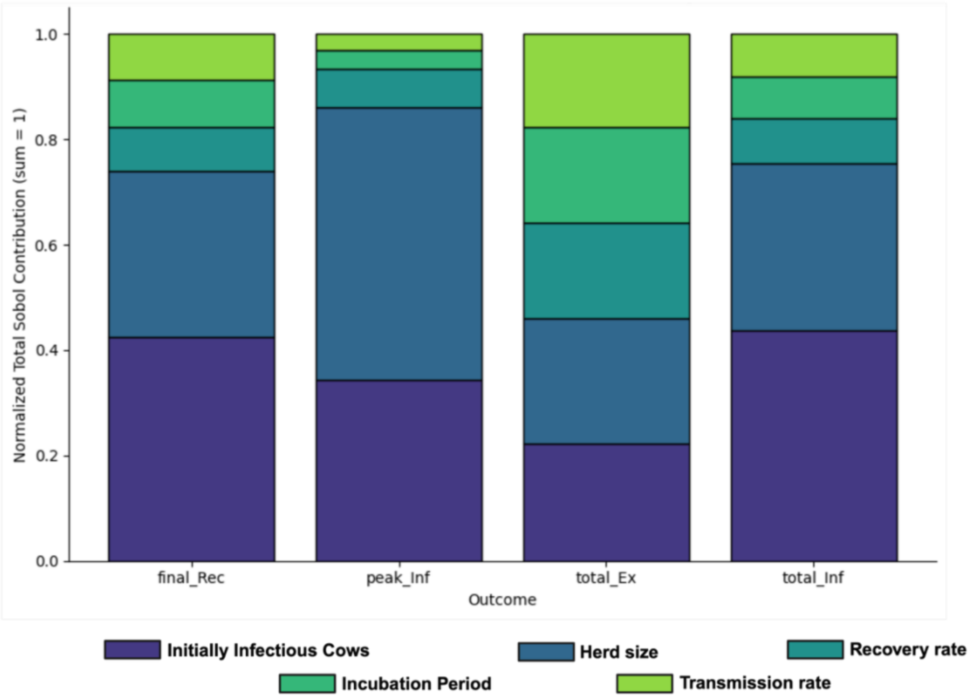

### Estimation of the transmission dynamics of H5N1 HPAI outbreak in a dairy herd using a modeling approach

**Pranav S. Kulkarni , Sharif S. Aly ,  Deniece R. Williams , Wagdy R. ElAshmawy ,**

**Pranav S. Pandit**

**Figure S4. Normalized contributions of input parameters to variance in outcome variables using Sobol method of global sensitivity. Abbrev. in figure: final_rec = Final Recovered, peak_Inf = Peak Infectious count, total_Ex = Total/ Cumulative Exposed count, total_Inf = Total/ cumulative Infectious count.**
