## Supplementary Files ODD Protocol for "Estimation of the transmission dynamics of H5N1 HPAI outbreak in a dairy herd using a modeling approach"

### Supplementary Material: ODD protocol for Model description

### Estimation of the transmission dynamics of H5N1 HPAI outbreak in a dairy herd using a modeling approach

**Pranav S. Kulkarni , Sharif S. Aly , Deniece R. Williams , Wagdy R. ElAshmawy , Pranav S. Pandit**

#### Overview

##### Purpose and pattern

The model was designed to simulate intra-herd outbreak of HPAI infection in dairy herd of Central Valley, California. The model was fitted with a parameter optimization protocol to match the temporal pattern of case data recorded on the dairy farms during September-October 2024.

##### Agents (entities) & Variables

**Table S1. Hierarchical structure of agents in Intra-farm HPAI outbreak model**

| Agent Type | Description | Level |
|---|---|---|
| Cow Agent | Represents individual cows with unique attributes; Dwells/moves between pens; epidemiological state tracking | 1 |
| Pen Agent | Represents functional pens that house cows with specific functions on the dairy farm | 2 |
| Farm Agent | Represents the whole farm, which contains all pens and cows; manages higher-level dynamics and herd decisions, pen/cow organization. | 3 |

**Table S2. Attributes of hierarchical agent-based simulation model of dairy herds**

| Attribute Name | Description |
|---|---|
| **Cow Agent** | |
| Cow ID | The unique identification number assigned to each cow upon entry into the herd (either as a calf or as a springer) |
| Current Pen | The current pen in which the cow agent is situated |
| Dwell time remaining | Remaining time duration for which the cow agent will remain in the current pen agent |
| Alive/ Culled | Binary status, whether the cow agent is active or has been culled (removed from further simulation) |
| Lactation Number | The number of lactations/ calves the cow agent has birthed |
| Days in Milk | The number of days cow agent dwells in milking pens |
| Epidemiological state | Four possible states: Susceptible (S), Exposed (E), Infectious (I) or Recovered (R) |
| Pen history | Container for storing records of which pens the cow agent has dwelled in, historically |
| **Pen Agent** | |
| Pen ID | Unique identification for Pens |
| Dwell time allocator | The module which tracks the dwell time assigned to each cow agent in the pen until the cow agent moves to the next pen. (Assigned stochastically when the cow agent moves into this pen agent.) |
| Transition rule – Next Pen | The module that tracks where each cow agent will move next to when the dwell time remaining reaches 0. |

### Estimation of the transmission dynamics of H5N1 HPAI outbreak in a dairy herd using a modeling approach

**Pranav S. Kulkarni , Sharif S. Aly , Deniece R. Williams , Wagdy R. ElAshmawy , Pranav S. Pandit**

| | |
|---|---|
| Susceptible cows | The counter for cow agents within the pen with epidemiological status of S |
| Exposed cows | The counter for cow agents within the pen with epidemiological status of E |
| Infectious cows | The counter for cow agents within the pen with epidemiological status of I |
| Recovered cows | The counter for cow agents within the pen with epidemiological status of R |
| **Farm Agent** | |
| Masked Pens | Pen IDs which need to be masked from epidemiological module. For example, open farms don't have calf pens on-site, hence, the calf pens are masked in SEIR dynamics. |
| Cow count | The counter for cow agents across all pens pooled to give the herd size in each time step |
| Culling candidates | In each time-step, the candidate cows for culling are selected randomly |
| History of Transitions | The module that stores all the histories of the cows on farm including culled cows |
| Day count | The counter for time steps |
| Calf count | The counter for newly initialized cow agents (calves) within the farm. Also assigns them unique Cow ID. |
| Susceptible cows | The herd-level counter for cow agents with epidemiological status of S |
| Exposed cows | The herd-level counter for cow agents with epidemiological status of E |
| Infectious cows | The herd-level counter for cow agents with epidemiological status of I |
| Recovered cows | The herd-level counter for cow agents with epidemiological status of R |

**Table S3. All the variables in the simulation module**

| Name | Description | Notation | Type |
|---|---|---|---|
| Pen dwell times | Represents the range of time spent by each cow agent in a particular pen agent. These time ranges are sourced from prior study[1] | $t_i$ | Deterministic |
| Culling rate | Represents a binomial sampling of culling probability of the farm. | $\mu_{annual}$ (%/ cows-days) | Deterministic |
| Transmission rate | Rate of transmission between infectious cows and susceptible cows upon contact. | $\beta$ | Stochastic |

[1] Konboon M, Bani-Yaghoub M, Pithua PO, Rhee N, Aly SS. A nested compartmental model to assess the efficacy of paratuberculosis control measures on U.S. dairy farms. PLoS One. 2018 Oct 2;13(10):e0203190. doi: 10.1371/journal.pone.0203190.

### Estimation of the transmission dynamics of H5N1 HPAI outbreak in a dairy herd using a modeling approach

**Pranav S. Kulkarni , Sharif S. Aly , Deniece R. Williams , Wagdy R. ElAshmawy , Pranav S. Pandit**

| | | | |
|---|---|---|---|
| Recovery rate | The rate of recovery of infectious cows wherein they are no longer infectious and able to transmit the infection. By definition, it is the inverse of infectious period for each cow ($T_I$). | $\gamma$ $\left(\frac{1}{T_I}\right)$ | Stochastic |
| Incubation period | The latent time between which a susceptible cow is successfully infected but has not become infectious. | $\delta$ | Deterministic |

##### Scales and duration

**Burn-in simulation**: Pre-simulation for setting up the dairy herd. Total duration is 3650 days (representing 10 years with 365 days in each year).
**Main simulation**: Variable duration depending on the cases reported by each of the modeled dairy farms. The range was between 17 and 53 days. For sensitivity analysis, this duration was increased to 100 days.
**Time step**: The simulation model progressed with a discrete time step of 1 day. Each of the movements, variables and probabilities were scaled to daily rates and proportions.

##### Process Overview and Scheduling

*Process*

The following pseudocode explains the process of simulation.
Pseudocode for Cow-Pen-Farm Simulation

- Initialize Farm agents
    - Create Pen agents for each pen definition (from some configuration/data).
    - Store each Pen agent in a pens dictionary in the Farm.

1. Add Initial Cow agents
    - For each cow configuration:
        - Create a Cow agent in the starting Pen, lactation state, and health/infection state.
        - Add Cow to both the Farm agent and the target Pen agent's cow list.
        - Initialize dwell start day for the cow agent.

2. Add Initial Infected Cows, if any.
    - If NOT burn-in simulation, at the start of the simulation:
        - Add a certain number of “Infectious” Cow agents, assign to random pen agents
    - Else,
        - Do not add any infectious Cow agents
3. Loop Over Each Simulation Day

### Estimation of the transmission dynamics of H5N1 HPAI outbreak in a dairy herd using a modeling approach

**Pranav S. Kulkarni , Sharif S. Aly , Deniece R. Williams , Wagdy R. ElAshmawy , Pranav S. Pandit**

- For each day in the simulation period:
    - For each Pen agent:
        - Identify Susceptible and Infectious cow agents.
        - Compute transmission probability and expose susceptible cow agents accordingly.
    - For each Cow agent:
        - Skip if agent is not alive.
        - Transition exposed to infectious, or infectious to recovered, based on infection time, incubation period, and recovery rates.
        - Call Cow.step(day, pens, farm), which moves the agent or progresses states as needed.
            - If time remaining in pen agent is 0, determine next pen agent or process special conditions (e.g., group dwell extension to match days in lactation constraint).
            - Move agent to the next pen agent, update pen agent's cow lists, and cow agent's dwell history.
            - If calving occurs (based on pen/lactation rules), reset lactation, add calf (new agent).
            - Farm agent adds new agents (calves) to the initial pen if calving occurred.
            - Apply mortality (randomly remove a percentage of living agents).

4. Record History and Aggregate Output Data
    - At each day, for each cow/pen agent:
        - Record relevant history data (state, pen, lactation) IF past burn-in.
        - Update farm-wide and pen-wide records for future analysis.

5. After Simulation
    - Aggregate and report on:
        - Cow histories.
        - Pen counts/time series.
        - Movements data.
        - Calving intervals and other summary statistics.

*Scheduling*

The following algorithm showcases the specific order by which each process is done.
Step 1:

```
1. INPUTS: load pen structure, simulation parameters (days,
epidemiological rates and parameters, etc.)

2. INITIALIZATION:
    - For each pen in configuration:
        - Create Pen object with ID, dwell min/max, next pen ID.
        - Add Pen to Farm's pens dictionary.
```

```
    - For each initial cow to add:
        - Create Cow in specified Pen and state.
        - Add Cow to Farm and Pen.
        - Set initial dwell start day.
    - If infectious cows required for simulation:
        - Add infectious cows to infectious pen.

3. SIMULATION LOOP:
For absolute_day from burn_in_days+1 to burn_in_days+ndays:
    a. For each cow in Farm.cows:
        i. If cow not alive, skip.
        ii. If cow.state == 'Exposed' and incubation period elapsed:
            - Set cow.state to 'Infectious'
            - Assign recovery day based on geometric sampling.
        iii. If cow.state == 'Infectious' and recovery day reached:
            - Set cow.state to 'Recovered'
            - Clear recovery_day
        iv. Record cow's current state/history if needed.
        v. Call cow.step(absolute_day, pens, farm):
            - Decrement time_remaining, increment dwell in current
lactation.
            - If time for new pen:
                - Determine next pen.
                - Check for group dwell extensions, update dwell time if
needed.
                - If moving to calving pen and lactation applies:
                    - Increment lactation, record calving interval, reset
calving-related counters, add calf.
                - Move cow: remove from old pen, add to new, update
time_remaining and dwell start day.
    b. For each pen in Farm:
        i. Identify living Infectious and Susceptible cows.
        ii. Compute exposure probability:
            - If both groups present, randomly select susceptible cows to
expose.
            - Set state and infection time, as needed.
    c. Apply daily mortality:
        - For all alive cows, select cows to be randomly removed based on
annual rate converted to probability.
        - Remove dead cows from pens.
    d. For each calf born:
        - Add new cow in starting pen.
    e. Record daily stats (cow histories, pen counts, movements).

4. POST-SIMULATION:
    - Aggregate and export histories (cow, pen, calving, dwell times,
etc.)
    - Analyze results as needed.
```

### Estimation of the transmission dynamics of H5N1 HPAI outbreak in a dairy herd using a modeling approach

**Pranav S. Kulkarni , Sharif S. Aly , Deniece R. Williams , Wagdy R. ElAshmawy , Pranav S. Pandit**

#### Design Concept

##### Basic Principle

To simulate the outbreak of HPAI, we developed a hierarchical agent-based simulation model. This model not only simulates the transmission of HPAI between infectious and susceptible cows but also tracks the cow movements between different pens on the farm. This methodology was developed using logical steps mentioned in Tedeschi (2023) [2]. The hierarchy of this model differs slightly from the traditionally used agent-based models (see review by Kaniyamattam and Tedeschi, 2023 [3]) wherein the pens are not treated as environment but as aggregating agents to allow for stochasticity in the dwelling times for each cow. In the same fashion, farm is treated as the higher-level agent rather than environment to allow for future expansion in modeling inter-farm HPAI transmission.

##### Emergence

Our simulation model will have emergent epidemiologic properties such as herd-level SEIR dynamics of interacting cow agents. We also expected the model to capture the emergent irregularities (heterogenous mixing) and stochasticity of agent-agent interactions within herds due to movement of cows between pens.

##### Adaptation

The only adaptative property that the agents in our model show are the movement of cows after the assigned pen dwelling time has ended. Each cow is assigned a random dwelling time that lies within the practical bounds of the ranges derived from standard dairy practices. Once assigned, these times are fixed and for each time step, the cow agents retain information on the time remaining and the pen agents retain information on the next pen agent the cow will move to deterministically. The farm agents control and handle all initializations including pen and cow agents, track the history and records and enforce the culling agenda by selecting candidate cow agents to be culled in each time step.

##### Objectives

Cow agents have multiple actionable objectives such as move between pens, remain alive or get culled, track lactations and days in milk. However, the objectives are not mapped onto any reward function given the nature of the model and the purpose.

---

[2] Tedeschi, L. O. 2023. Review: the prevailing mathematical modelling classifications and paradigms to support the advancement of sustainable animal production. Animal 100813. doi: 10.1016/j.animal.2023.100813

[3] Kaniyamattam K., Tedeschi L. O. 2023. ASAS-NANP symposium: mathematical modeling in animal nutrition: agent-based modeling for livestock systems: the mechanics of development and application. Journal of Animal Science 101, skad321. doi: 10.1093/jas/skad321

### Estimation of the transmission dynamics of H5N1 HPAI outbreak in a dairy herd using a modeling approach

**Pranav S. Kulkarni , Sharif S. Aly , Deniece R. Williams , Wagdy R. ElAshmawy , Pranav S. Pandit**

Pen agents have objectives such as track and hold cow agents, identify the next pen for each cow agent (update membership). For milking pens, a specialized objective of extending dwelling time is attached so that cows complete the biological milking cycle of at least 305 days.
Farm agents have objectives such as track all pen and cow agents and record history.

##### Sensing and Learning

Our simulation model is set up on a rule-based operating procedure and does not have learning or sensing capabilities.

##### Interaction

The interactions between cow agents are primarily driven by being in the same pen at the same time step. This is the primary driver of transmission of HPAI from infectious to susceptible cow agents. Cow agents in different pens do not interact and therefore do not transmit infection. Pen agents interact through cows wherein each pen determines the next logical pen for each cow inside the pen to move into after the dwelling time in the current pen has ended.

##### Stochasticity

There are two stochastic components to our model: the dairy operational stochasticity and the epidemiological stochasticity. In the dairy operational part, the main stochastic elements are the selection of candidate cow agents to be culled in each time step and the bounded random assignment of dwelling times for each cow in each pen. In the epidemiological part, the main stochastic elements are probability of exposure for susceptible cows when they interact with infectious cows from the same pen (based on transmission rate parameter) and the recovery rate for each infectious cow. For specific sampling distributions refer to equations in the main Methods section of the manuscript.

##### Collectives

Both Pen and Farm agents are nested aggregating agents that form collectives of cow agents on pen and herd level respectively.

##### Observation

Each cow agent records epidemiological status, history of its movements between different pens, lactation number (and number of calves born), days in milk for current lactation, culling status. Each pen agent collects the current count of cows in the pen, the tracking of the next pen agent for each cow, assigned dwell time for each cow coming into the pen and the counts for SEIR cows. Each farm agent collects all the history of movements of the existing and previously culled cows, the current candidates for culling, the total herd size (cow agent count with alive status vs culled status), number of pens and pens to be masked in epidemiological simulations.

### Details

#### Initialization and Input data

**Table S4. Initial inputs and conditions for hierarchical agent-based model**

| Inputs | Units | Values (Range) |
|---|---|---|
| Number of simulations | - | Variable (100-1000) |
| Number of pens | - | 15 |
| Initialization pens | - | Pre-weaning, Springer pens |
| Target herd size | Animal units | 800, 500, 2500, 2500 |
| **Burn-in simulation** | | |
| Initial Number of cows | Animal units | § 300, 200, 900, 900 |
| Initial Number of calves | Animal units | § 300, 200, 900, 900 |
| Duration | Days | 3650 |
| Culling rate (herd level) | % | 30% |
| Initially infectious cows | Animal units | 0 |
| Transmission rate | - | 0 |
| Recovery rate | - | 0 |
| Incubation period | Days | 0 |
| **Main simulation** | | |
| Initial number of cows | Animal units | Burn-in herd size |
| Initial number of calves | Animal units | Burn-in calves |
| Duration | Days | § 33, 53, 23, 18 |
| Culling rate (herd level) | % | 30% |
| Initially infectious cows | Animal units | Variable (1-50) |
| Transmission rate | - | Variable (0.01-0.99) |
| Recovery rate | - | Variable (0.01-0.99) |
| Incubation period | Days | Variable (1-10) |

**§ Inputs for specific farms: Farm1, Farm2, Farm3, Farm4 respectively**

**Table S5. Dwelling time range for each pen and next pen for transition after end of dwelling time.**

| Current Pen | Minimum duration (days) | Maximum Duration (days) | Next Pen | Group |
|---|---|---|---|---|
| Pre-weaning (Pen 1) | 60 | 80 | Post-weaned (Pen 2) | Calf/ Young cow |
| Post-weaned (Pen 2) | 396 | 458 | Breeding (Pen 3) | Calf/ Young cow |
| Breeding (Pen 3) | 60 | 100 | Pregnant (Pen 4) | Calf/ Young cow |
| Pregnant (Pen 4) | 234 | 264 | Springer (Pen 5) | Calf/ Young cow |
| Springer (Pen 5) | 18 | 32 | Hospital (Pen 15) | Primiparous |

| | | | | |
|---|---|---|---|---|
| Fresh Primiparous (Pen 6) | 5 | 30 | High milking Primiparous (Pen 8) | Primiparous |
| Fresh Multiparous (Pen 7) | 30 | 90 | High milking Multiparous (Pen 10) | Multiparous |
| High milking Primiparous (Pen 8) | 180 | 200 | Low milking Primiparous (Pen 9) | Primiparous |
| Low milking Primiparous (Pen 9) | 100 | 160 | Drying (Pen 12) | Primiparous |
| High milking Multiparous (Pen 10) | 90 | 180 | Low milking multiparous (Pen 11) | Multiparous |
| Low milking multiparous (Pen 11) | 200 | 360 | Drying (Pen 12) | Multiparous |
| Drying (Pen 12) | 48 | 66 | Close-up (Pen 13) | Multiparous |
| Close-up (Pen 13) | 1 | 7 | Calving (Pen 14) | Multiparous |
| Calving (Pen 14) | 1 | 7 | Hospital (Pen 15) | Multiparous |
| Hospital (Pen 15) | 2 | 2 | Fresh Primiparous or Fresh Multiparous (Pens 6 or 7) | - |

#### Sub-models

The sub-models used are as follows:

*Cow agent*

1. **Step function**: This function accounts for the dwelling time remaining in the current pen and decreases it by 1 for each time step. If the current pen is hospital pen (Pen 15), it assesses the lactation number of the cow to direct the cow into either Fresh Primiparous (Pen 6) or Fresh Multiparous (Pen 7) pens respectively for lactation = 1 or > 1. After the cow agent enters the hospital pen, this function indicates to the farm agent that a calving has occurred, and a new cow agent (calf) is initialized. Further, the function puts constraint of 305 days on the total dwelling time of milking pens to avoid biologically unfeasible milking durations.
2. **Calculate infectious period**: This function calculates the time remaining in the infectious state and decreases it by 1 for each time step.
3. **Recording functions**: These are a group of functions that record the movement data, change in epidemiological status and directs the prior records to pen and farm agents.

*Pen agent*

4. **Add cow function**: This function allows and records entry of every new cow agent and assigns them the dwelling time within the dwelling time range.
5. **Remove cow function**: This function allows movement of cow from this pen agent when dwell time remaining is equal to 0.
6. **Get Next Pen ID function**: This function assigns the next pen agent into which a cow agent will move when the dwell time is equal to 0. (Exception: Pen 15/ Hospital Pen, handled within the cow agent).

### Estimation of the transmission dynamics of H5N1 HPAI outbreak in a dairy herd using a modeling approach

**Pranav S. Kulkarni , Sharif S. Aly , Deniece R. Williams , Wagdy R. ElAshmawy , Pranav S. Pandit**

*Farm agent*

1. **Add cow function**: This function initializes new cows (initially introduced calves or springers or calves born during simulation) and assigns a unique Cow ID to the initialized agents.
2. **Step day function**: The main sub-model that simulates one day time step. This function performs the following actions:
   a. Counts number of cow agents to be culled (removed from further simulation). Selects candidate cows for culling randomly from the active cows by sampling active cow agents using the culling rate
   b. Changes epidemiological states for S, E, I and R cows based on transmission rate, recovery rate and end of infectious period (calculated by Calculate infectious period function in cow agent). Also tracks the number of infectious cows and susceptible cows in each pen to perform these dynamics which are later summed up on herd-level.
   c. Tracks number of new calves born during the simulation
3. **Run simulation function**: Initializes simulation runs (either burn-in simulation or main simulation which is user-input defined) and creates containers for recording data.
4. **Initiate infectious cows**: Introduce new cows with epidemiological state set to “Infectious” to random pens (or specific pens such as High Milking pen/ Pen 8, based on user-input). Also assigns their infectious period based on recovery rate.
5. **Recording functions**: Group of recording sub-models which record the movements, appends history, collects lactation data of cows, logs all completed dwell time for debugging in case of errors
